# Ultra high-throughput whole-genome methylation sequencing reveals trajectories in precancerous polyps to early colorectal adenocarcinoma

**DOI:** 10.1101/2022.05.30.494076

**Authors:** Hayan Lee, Gat Krieger, Tyson Clark, Aziz Khan, Casey Ryan Hanson, Yizhou Zhu, Nasim Bararpour, Aaron M. Horning, Edward D. Esplin, Stephanie Nevins, Annika K. Weimer, Eti Meiri, Shlomit Gilad, Sima Benjamin, Danit Lebanony, Nika Iremadze, Florian Oberstrass, Ariel Jaimovich, William Greenleaf, James M. Ford, Doron Lipson, Zohar Shipony, Michael P. Snyder

**Author notes:** Contributed equally to this work (Hayan Lee, Gat Krieger).

## Abstract

Aberrant shifts in DNA methylation have long been regarded as an early marker for cancer onset and progression. To chart DNA methylation changes that occur during the transformation from normal healthy colon tissue to malignant colorectal cancer (CRC), we collected over 50 samples from 15 familial adenomatous polyposis (FAP) and non-FAP colorectal cancer patients, and generated 30-70x whole-genome methylation sequencing (WGMS) runs via the novel Ultima Genomics ultra high-throughput sequencing platform. We observed changes in DNA methylation that occur early in the malignant transformation process, in gene promoters and in distal regulatory elements. Among these changes are events of hyper-methylation which are associated with a bivalent “poised” chromatin state at promoters and are CRC-specific. Distal enhancers show nonlinear dynamics, lose methylation in the progression from normal mucosa to dysplastic polyps but regain methylation in the adenocarcinoma state. Enhancers that gain chromatin accessibility in the adenocarcinoma state and are enriched with HOX transcription factor binding sites, a marker of developmental genes. This work demonstrates the feasibility of generating large high quality WGMS data using the Ultima Genomics platform and provides the first detailed view of methylation dynamics during CRC formation and progression in a model case.

## Introduction

CRC is the third most diagnosed cancer in men and second in women and accounts for 8% of all cancer deaths^1^. Recent improvements in colorectal cancer diagnosis and therapeutic strategies have increased CRC patient survival time while the mortality rate of CRC remains high^2^.

The molecular pathway causing CRC is the conventional adenoma–carcinoma pathway^3^. The pre-malignant to malignant progression of sporadic adenoma to carcinoma has been described for CRC malignancies and occurs in a stepwise fashion^3,4^. The initiating step in 80-90% of colorectal tumors is the loss of adenomatous polyposis coli (APC) gene resulting in β-catenin stabilization and increased WNT signaling^5^. Subsequent mutations in other cancer driver genes such as KRAS, TP53, and SMAD4 result in the transformation to carcinoma. Familial adenomatous polyposis (FAP) is an autosomal dominant inherited disorder caused by a germline mutation in the APC gene. The disease is characterized by the formation of numerous adenomatous polyps mainly in the epithelium of the large intestine, typically arising at teenage years^6,7^. This leads to a very high likelihood of malignant transformation in at least one of these polyps by the fifth decade, 100% lifetime risk of CRC and is only preventable by prophylactic removal of the entire colon (colectomy)^7^. The presence of benign and dysplastic polyps in FAP patients is thought to reflect the earliest steps in CRC formation, and as such presents a valuable and unique system for dissecting the early events in CRC tumor initiation and progression in the same initial genetic background.

DNA methylation is one of the most widely studied epigenetic modifications. Changes in DNA methylation correlate with changes in gene expression and cell state^8,9^, and have been observed in virtually every cancer type^10^. Cancer cells typically exhibit global hypo-methylation while specific loci such as CpG islands are usually hypermethylated^11–13^. These alterations occur very early in the carcinogenesis process, in many cases before malignant transformation, and increase with tumor progression^14,15^. DNA methylation changes specific to CRC have been previously reported in many genomic regions including gene promoters, LINE1 repeat elements and at regulatory regions bound by the Polycomb group protein complex^16,17^. A distinguished type of regulatory regions that show increased methylation in cancer, are bivalent promoters and enhancers (also called poised promoters/enhancers). These regions are enriched with activating and repressing chromatin modifications that co-occur at the same genomic regions and are pre-loaded with poised RNA polymerase II to prepare genes for rapid activation. Poised promoters are typical for developmental genes and were suggested to “safeguard differentiation” and thus their malfunction is expected to have a harmful impact on the cell^18–20^.

In recent decades, the availability of next generation sequencing technologies has profoundly improved our understanding of the molecular basis of human disease and enabled data-informed drug design, personalized disease treatment and improved monitoring of disease progression^21,22^. However, in recent years, the decrease in sequencing cost has plateaued and has been a limiting factor in both the number of samples interrogated and in the scope of data collected per sample, especially for individual laboratories^23^. To date, most studies, including large cancer genome profiling studies such as The Cancer Genome Atlas (TCGA)^21^ have opted to measure genomic methylation status using targeted approaches like arrays rather than genome wide approaches due to the high cost of sequencing associated with whole genome methylation sequencing (WGMS) at relevant coverage.

The Ultima Genomics (UG) sequencing platform utilizes a new sequencing architecture that combines an open flow cell design on a circular wafer with large surface area, utilizing rotational reagent delivery, optical end-point detection, and mostly natural nucleotides without reversible terminators. This platform enables sequencing billions of reads with high base accuracy (Q30 > 85%), at significantly reduced cost versus conventional sequencing platforms^24^, thereby allowing efficient generation of comprehensive WGMS data. In this study we used the UG platform to sequence 44 WGMS samples along the FAP tumor progression from normal mucosa to adenocarcinoma at high coverage (30-60X). This broad WGMS dataset allowed us to better monitor the transformation happening at the early stages of cancer development and discover thousands of methylation changes that occur at the transition from the normal mucosa stage to both the benign and dysplastic stages before adenocarcinoma formation. Using WGMS we were also able to assess hundreds of thousands of CpG methylation changes that occur at distal regulatory elements, most of which were not previously measured by enrichment and array-based assays. In addition to presenting this novel method and results, the data serve as a valuable scientific resource for probing early events associated with CRC.

## Results

### Ultima Genomics whole genome methylation

To compare WGMS data generated by the UG sequencer to WGMS methods we sequenced the standard reference HG001, HG002 and HG005 cell-lines using both EM-seq^25^ and Bisulfite^26^ conversion methods and compared our data with the recently published Genome in a Bottle (GIAB) EpiQC data^27^. Using the UG sequencer we sequenced an average of 922 million reads per sample for each of the methods with two to five replicates (Supplementary table 1). We first examined general mapping statistics such as read mappability rate, duplication rate and the percent of Lambda (fully unmethylated) and pUC19 (fully methylated) control DNA spike-ins. Overall, UG datasets had high mappability rates (95% to 99%) and low duplication rates (15% to 30%; Supplementary Fig 1a-b). The internal controls for methylation (Lambda and pUC19) showed high conversion efficiency both for EM-seq (98.5%) and Bisulfite-seq (97%) (Fig 1a-b) and C to T conversion rates at unmethylated CpGs were uniform along the read (Supplementary Fig 1c).

**Figure 1:**
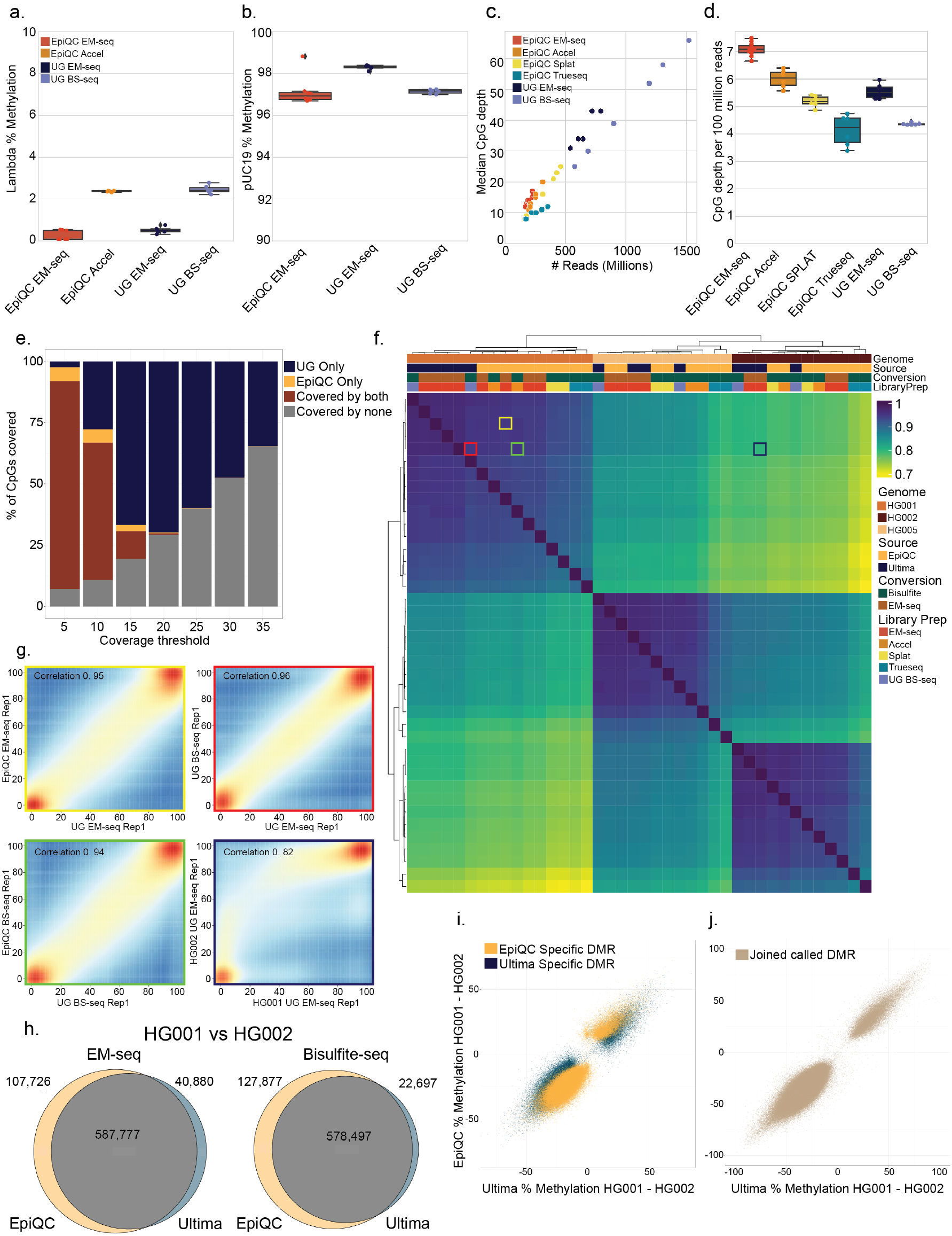
Comparison between Ultime Genomics sequencing of to EpiQC EpiQC reference set. (a-b) Average methylation levels (y-axis) at control sets, Lambda non-methylated DNA and pUC19 fully-methylated DNA in UG/EpiQC EM-seq (enzymatic conversion) libraries, in EpiQC Accel-NGS (bisulfite conversion) libraries and in UG xGen (bisulfite conversion) libraries. (c) CpG coverage in UG and EpiQC data, median CpG depth (x-axis) vs. millions of sequenced reads (y-axis). (d) CpG median coverage normalized to 100 million reads at all data sets. (e) Fraction of CpGs out of hg38 CpGs (29M) that are covered by by at least two replicates in none (gray), both (brown) or each of the two platforms (yellow and blue, EpiQC and UG respectively). (f) Correlation of whole genome methylation sequencing between cell lines in platforms. Heatmap shows the Pearson correlation coefficients between each two samples coming from different sources, assays and platforms. Top row denotes the source of the sample(orange, brown and yellow for HG001, HG002 and HG005 respectively). Second row denotes the two platforms (yellow and blue for EpiQC and Ultima Genomics, respectively). Third row denotes the two conversion methods (brown and green for EM-seq enzymatic conversion and bisulfite conversion, respectively). Fourth row denotes the library prep methods (red, orange, yellow, green and blue for EM-seq, Accel-NGS, Splat, Trueseq and xGen, respectively). (g) Examples of four pairwise correlation plots taken from Fig 1f (border color of each image is marked on the correlation map) (h-j) Differentially methylated regions (DMRs) are detected similarly in UG and EpiQC platforms. (h) DMRs between HG001 and HG002 genomes were called in both platforms. Venn diagrams show the overlap in DMRs between the two platforms, using two replicates from EM-seq (left) and two replicates from Bisulfite conversion (right) methods, (i) Platform-specific DMRs show a similar change in methylation. Shown is the observed methylation delta between each two replicates of HG001 and HG002 genomes in UG (x-axis) and in EpiQC (y-axis), color of the points encodes for platform-specificity (yellow and blue for EpiQC specific regions and UG specific regions respectively). (j) Observed methylation delta in methylation between each two replicates of HG001 and HG002 genomes in UG (x-axis) and in EpiQC (y-axis) in jointly-called DMRs.

We next examined the genomic coverage of CpG sites achieved by the UG in comparison to EpiQC data. UG achieved comparable coverage statistics to EpiQC, with median CpG depth demonstrating a linear increase in depth as function of numbers of reads (Fig 1c). At the read level UG achieved between 4.5 and 5.5 average CpG depth per 100 million reads vs 4-7 by EpiQC (Fig 1d, note that UG reads were generally slightly shorter than in EpiQC data, Supplementary table 1).

Importantly, the high UG coverage depth resulted in a much higher average coverage per CpG (Fig 1e). On average, we observed 70% of CpGs covered at over 20X leading to smaller deviations between replicates when compared to lower coverage (Supplementary Fig 1d).

To compare the methylation calls between the two sequencing platforms we first assessed the average methylation across all the samples sequenced (Supplementary Fig 1e). We observed high variability between the different library preparation methods (standard deviation between methods are 0.7, 1.2, 1.5 for HG001, HG002 and HG005), which is higher than the variability between the different platforms (standard deviation between platforms of EM-seq method are 0.57, 0.77, 0.62 for HG001, HG002 and HG005). Importantly, we observed that methylation patterns around the transcription start site (TSS) of all genes and across many different genomic features behaved similarly on both platforms (Supplementary Figs 1f & 2a).

To further test agreement between the different platforms we measured the correlation of methylation levels of all CpGs with >5X coverage in both EpiQC data and UG genomics data (24.5 M CpGs; Fig 1f). We observed a very high correlation (R = 0.94 to 0.96) when measuring methylation levels of the same cell line with the two technologies. Correlation between different conversion methods was similar to that of the correlation using the same conversion method (R = 0.94 to 0.96) between the two sequencing platforms (Fig 1g).

Finally, we evaluated differential analysis performed on the two sequencing platforms by identifying differentially methylated regions (DMRs) between HG001 and HG002. We found an 80% overlap between the detected DMRs of the different platforms for both EM-seq and Bisulfite-seq conversion (Fig 1h). Further inspection demonstrated that most differences in detected DMRs were explainable by marginal calls near the calling threshold (Fig1 i-j). Overall, these results indicate that the technical variation between UG and EpiQC is modest (R = 0.95) when compared to differences in biological samples from different sources (HG001 vs HG002; R = 0.82).

### DNA methylation changes with cancer development of Familial adenomatous polyposis patients

We next examined the ability of the UG platform for characterization of genome wide methylation changes during the early colon polyp formation using FAP samples. To this end, we collected blood and multiple fresh frozen samples of colon “normal” mucosa, polyps and an adenocarcinoma sample from nine FAP patients (Fig 2a; Table 2; Methods). We also collected fresh frozen samples from adenocarcinoma tissues from six CRC patients. Tissue slides of the normal mucosa and polyps were examined by a pathologist and defined into four categories of increasing severity: normal mucosa, benign, dysplastic and adenocarcinoma. DNA was extracted from these samples, processed using EM-seq, and sequenced to an average genomic coverage of ~50 X (25x to 112x) (Fig 2a and Supplementary Fig 3b).

**Figure 2:**
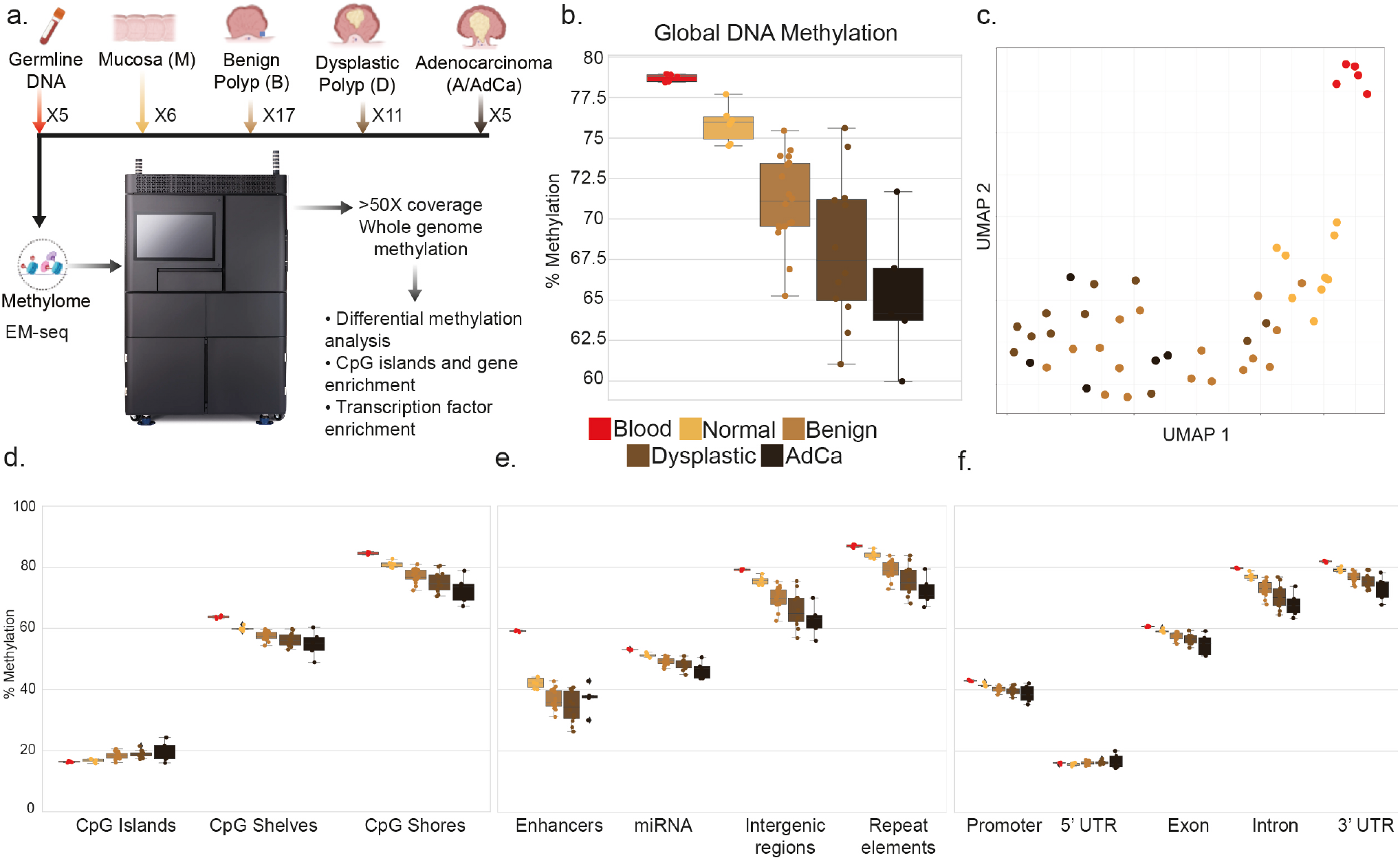
Familial adenomatous polyposis samples collected across different pathologist-defined states show changes in DNA methylation at malignant states trajectories. (a) Whole genome methylation sequencing at high coverage was done from four different pathologist-defined states: normal mucosa, benign, dysplastic and adenocarcinoma. Blood samples were collected from the same individuals for reference. (b) Average genome-wide DNA methylation levels at each of the pathologist-defined states(red, yellow, light brown, dark brown and black for blood, normal mucosa, benign, dysplastic and adenocarcinoma, respectively). (c) UMAP representation of WGMS samples done on CpGs where all samples have at least 10X coverage per CpG (colored same as in (b)) (d-f) Average DNA methylation levels of all disease states (colored same as in (b)) by different genomic features (x-axis).

To gain insight on the tissue composition of these samples we applied a computational method for methylation-based cell type deconvolution^28^(methods; Supplementary Fig 3c). This analysis revealed that three of the adenocarcinoma samples had a negligible colon fraction and one of them contained a high fraction of lung cells. Pathologist review on Hematoxylin and Eosin (H&E) slides taken from FFPE samples from some of the same tissues further validated the tissue composition estimation for 2 of 3 cases (Supplementary Fig 3d) including the observation of significant lung cell infiltration in one of the samples. To avoid the possible confounding effect of significant non-colon tissue in the samples we set a threshold of 35% colon fraction as the minimal requirement for downstream analysis, excluding six samples (3 AdCa, and 3 normal mucosa polyps) from further analysis.

Average genome wide methylation levels show a gradual decrease during tumor progression from a median of 75% genomic methylation in normal tissues to ~65% in adenocarcinoma samples (Fig 2b; Supplementary Fig 3a) in agreement with existing studies^11,29^. To visualize the change of methylation states during disease progression we projected the observed global methylation levels using the UMAP dimensionality reduction technique (Fig 2c). This embedding clearly differentiates the blood samples from the colon ones and suggests a gradual change that is largely consistent with pathological grading of the tissues. Pearson correlation of the global methylation levels supports similar observations (Supplementary Fig 3f).

To further examine whether this trend is consistent in different functional genomic regions we measured methylation changes in CpG Islands/Shores/Shelves (Shores: 2kb around the CpG islands, Shelves: 2k-4kb around the CpG island), repeat elements and other regulatory elements. (Fig 2d-f). In contrast to most genomic features, CpG islands exhibit a slight increase in methylation during cancer progression while the CpG shelves and shores show the same decrease as other regions (Fig 2e)^30^. 5’ UTR regions that typically contain many CpG islands also show a similar increase. Other genomic elements that show deviation from the global behavior are colon-specific regulatory regions and enhancers (Fig 2e and Supplementary Fig 3e); these regions lose methylation in the progression to dysplastic state and regain methylation from dysplastic to tumor state. In addition, we observe that the variability within genomic features increases with progression towards cancer (Fig 2d-f and Supplementary Fig 3e)

### Tens of thousands of differentially methylated regions (DMRs) are found between normal mucosa and adenocarcinoma states

To characterize genomic regions that change in methylation during tumor progression we performed a pairwise DMR^31^ test between normal mucosa (M) to the pathologist-defined sample groups (benign (B), dysplastic (D) and adeno (A), B-M, D-M, or A-M, respectively). Overall, we found 150,508 DMRs covering almost 3 million individual CpG sites, one of the largest such DMR maps ever reported. Partitioning DMRs to regions that lose methylation during disease progression (hypo) and regions that gain methylation during disease progression (hyper), we found that hypo-methylated DMRs are significantly more common than hyper-methylated DMRs (144,284 vs 6,224) and that while many hyper-methylated DMRs (30%) can already be seen in benign polyps, only a small fraction of hypo-methylated DMRs (10%) are detected first in the benign polyps (Fig3 a-b) suggesting different methylation dynamics over disease progression. As may be expected, the amplitude of methylation changes increases gradually from the benign DMRs (90% of the delta = 0.246) to the dysplastic and adenocarcinoma DMRs (90% of the delta = 0.318 and 0.348 respectively; Fig 3c). We next examined the differences in genomic features in each DMR group and found that CpG islands, CpG shelves, promoters and regulatory regions were overrepresented at hyper-methylated DMRs, while normally “inert” functional groups such as repeat elements and intergenic regions were over-represented at hypo-methylated sites (Fig 3d). This trend was consistent among the different disease states. An example for a cluster of hyper-methylated DMRs is the HOXA locus (Figure 3e). This genomic region of 35 kb has multiple DMRs that gradually elevate in methylation levels from normal mucosa to adenocarcinoma, and are associated with reduced accessibility as viewed by ATAC-seq data (Figure 3e). The expression of three HOXA genes is repressed in tumors as seen in TCGA data (Supplementary Fig 4a). HOX hyper-methylation was observed before in colorectal cancer as well as in multiple other cancers^32^.

**Figure 3:**
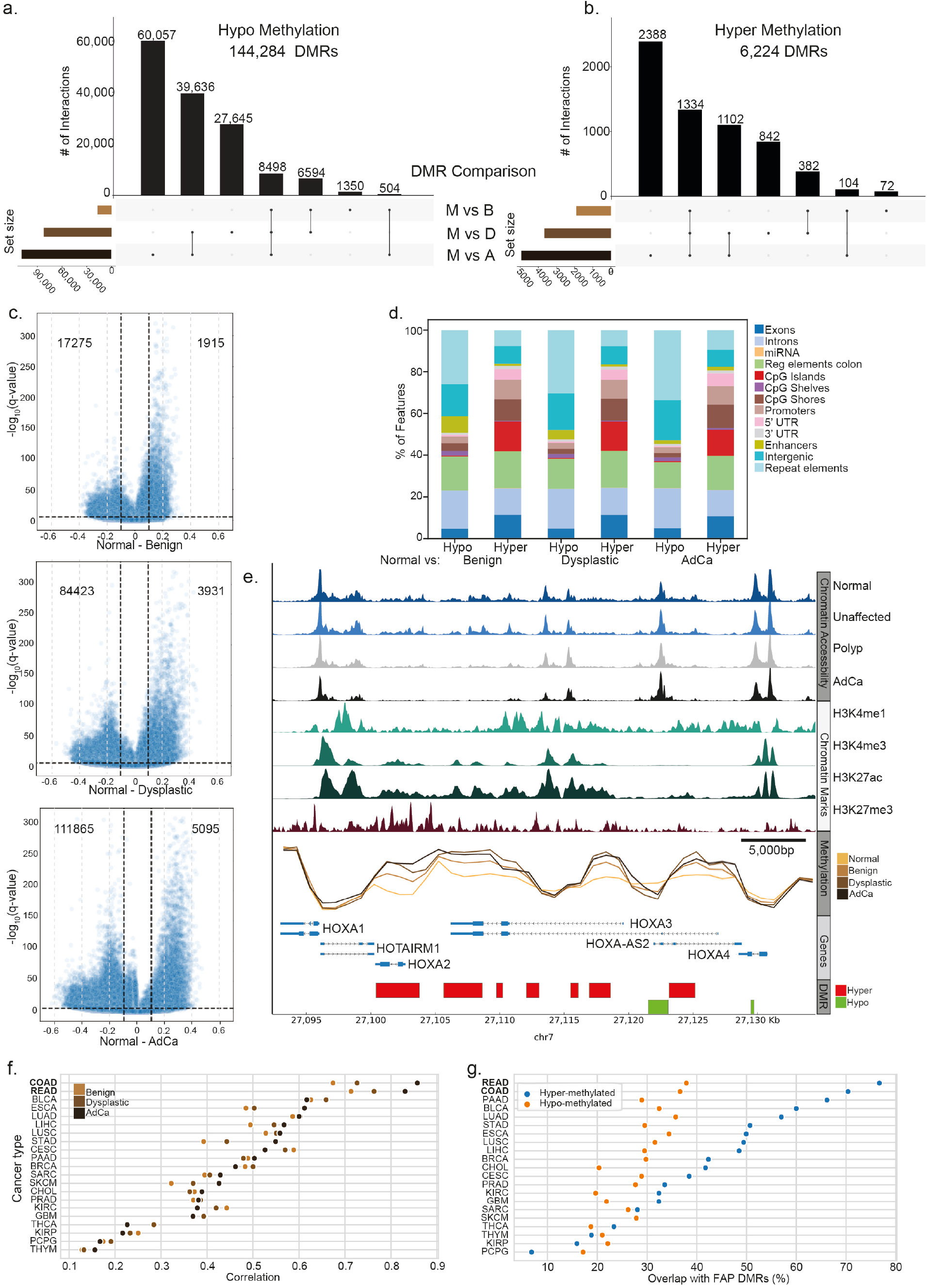
Differential methylated regions during progression from normal to malignant. (a-b) Intersect plot of pairwise comparison of normal vs benign, dysplastic and adenocarcinoma where horizontal bars show the size of each pairwise DMR set and the horizontal bars show the size of the respective overlap. (a) 144,284 Hypo-methylation DMRs between all pairwise tests. (b) 6224 Hyper methylation DMRs between all pairwise tests, 1334 of which occur early in the benign stage. (c) Volcano plots of differential methylation levels: shown are methylation differences compared to normal mucosa samples (x-axis) and q-value statistical significance (y-axis) of all pairwise comparisons. (d) Genomic annotation of different DMRs tests distinguished to hyper- and hypo-methylation. (e) *HOXA* cluster shows multiple DMR both at hyper and hypo methylation. (f-g) comparison to the TCGA dataset. (f) Change in methylation in CRC progression specifically matches colorectal adenocarcinoma (COAD) and rectal adenocarcinoma (READ). Shown is the Pearson correlation coefficient between the change in methylation from normal mucosa and disease states (color-coded) in our data and normal to tumor methylation changes in TCGA data of 450k arrays. Comparison includes 339,977 CpGs. (g) High fraction of the hyper-DMRs, but not the hypo-DMRs are differentially methylated in the TCGA dataset. Differential methylation in TCGA as determined by DNMIVD^33^ per CpG site and cancer type, and filtered to include methylation difference >15%, adjusted p-value <0.05. Presented is the percent intersection of the CpGs in DMRs we detect among differentially-methylated CpGs of TCGA.

We next wanted to test whether the gradual changes we observe in the different disease states are specific to CRC. To this end we compared the average per-CpG methylation change in each of the three disease states with the pan-cancer methylation changes in the same CpGs in hundreds of normal and tumor samples studied in TCGA^21^ (Fig 3f). As expected, the change in methylation in adenocarcinoma was mostly correlated with methylation changes in COAD (**Co**lon **adenocarcinoma**) and READ (**Re**ctum **ad**enocarcinoma) in TCGA (R > 0.8) whereas correlation levels with other cancer types were lower (R < 0.61). Of note, the change in methylation in the intermediate disease states (Benign, Dysplastic) were also specifically correlated with the same matched tumor types, and correlation increased with disease progression (Fig.3f). We also observed that hypo-methylated markers show a lower overlap (<40%) with TCGA data, suggesting that hypermethylated regions are more cancer type specific than hypo-methylated regions (Fig 3g).

Lastly we assessed how many of the DMRs that we detect are also found as differentially methylated in TCGA data^33^. Among the full set of DMRs we detected in all disease states (153,525 regions), only 15% are covered by Infinium HumanMethylation450 array (24,067 regions), occupying 71,462 array marker CpGs. Thus, WGMS provides a substantially broader view of dynamic DMRs and differentially methylated CpGs.

### Poised genes are hyper-methylated during cancer progression

We next explored the relationship between DNA methylation dynamics and chromatin architecture and their correlated changes during cancer progression. We first compared methylation levels observed in normal mucosa samples to the chromatin accessibility in the same regions in normal mucosa^34^. Grouping these regions according to predefined functional chromatin state clusters (ChromHMM^35^) captures different behaviors, where most regions show negative correlation between methylation levels and chromatin accessibility (Fig 4a). For example, active regions such as active and flanking TSS (groups 1-4) display high DNA accessibility levels and very low DNA methylation levels. Regions of strong and weak transcription (gene bodies, groups 5-6) have high DNA methylation levels and low accessibility levels. Regulatory regions (enhancers, groups 7-11) show an intermediate behavior where both accessibility and methylation are at intermediate levels. Bivalent enhancers and TSSs are the only group that demonstrates a different trend combining low accessibility levels and low methylation levels in normal colon mucosa tissues.

**Figure 4:**
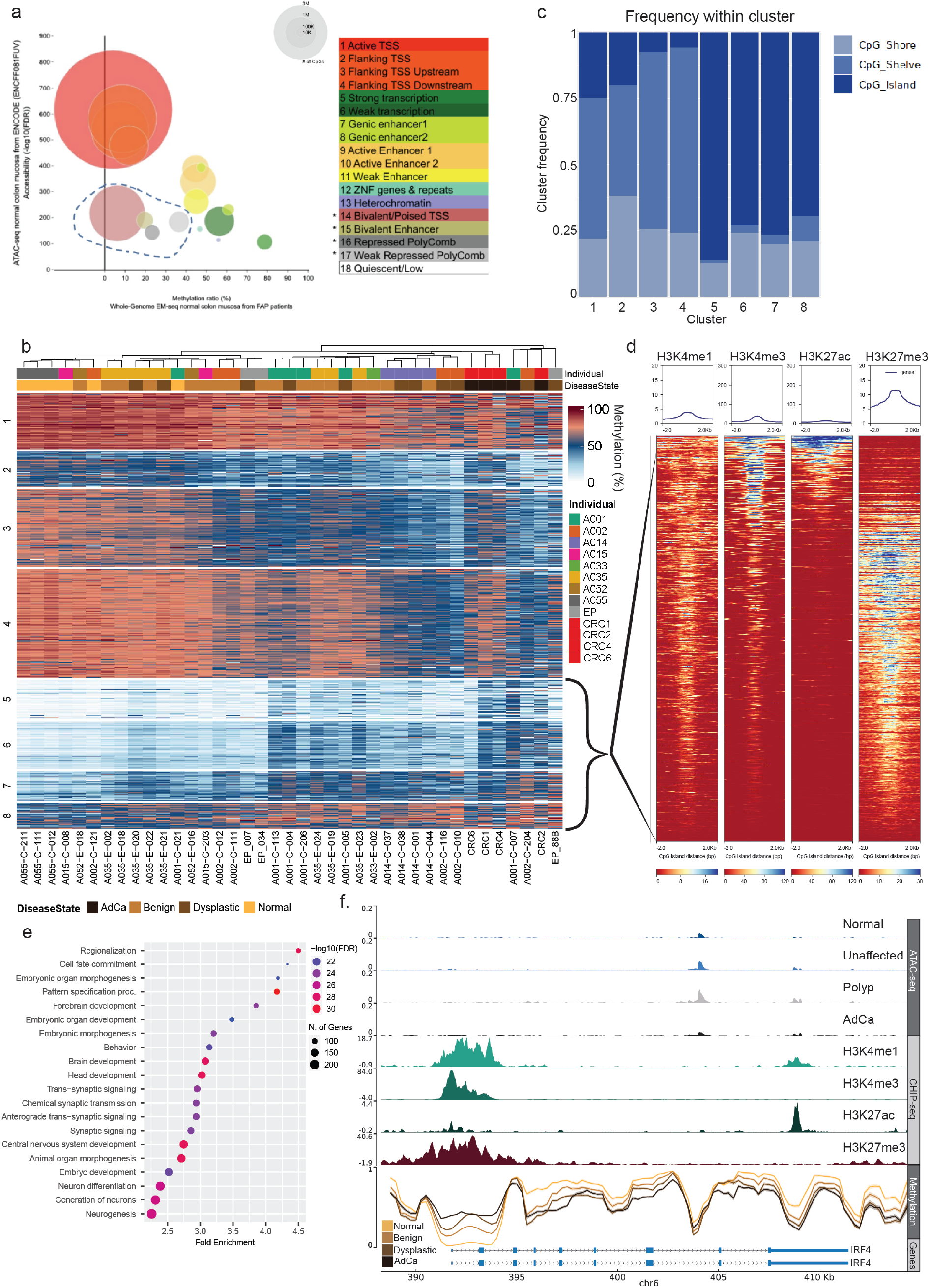
Poised genes are hyper-methylated during cancer progression. (a) DNA accessibility in normal colon samples (ENCODE) vs DNA methylation levels in normal mucosa (this study), grouped and colored by ChromHMM chromatin state. (b) K-Means (K=8) clustering of 11,323 variable CpG islands/shelves and shores (3825 CpG Islands, 4821 CpG shelves and 2677 CpG shores) across all FAP samples. Variability was determined as standard deviation >10 across all samples. First row denotes patient identity by color and second row denotes sample pathology label (yellow, light brown, dark brown and black for normal mucosa, benign, dysplastic and adenocarcinoma, respectively) (c) Fraction of annotation type (CpG island/shelves and shores) in each of the K-Means clusters. (d) Histone marks levels (H3K4me1/3, H3K27ac and H3K27me3) sigmoid colon (ENCODE, Normal colon) on clusters 5-8 from panel b. Each row shows the color coded histone mark level in the relevant genomic position centered around the CpG island/shore/shelf from low (red) to high (blue). Top panels show average marker level. (e) GO enrichment of genes associated with CpG islands of clusters 5-8 of panel b. Circle size correlates with the overlap between the GO term and the genes adjacent to the CpG islands in clusters 5-8. Circle colors maps the FDR corrected p-value for the enrichment (f) *IRF4* genomic track showing gradual increase in CpG methylation at the CpG islands next to the TSS and gradual decrease of CpG methylation at the gene body of *IRF4*. First four rows show the chromatin accessibility in the four pathology groups, next four rows show histone modification levels in normal mucosa and last panel shows DNA methylation level for the different groups as in panel b.

To better understand the relationship between chromatin and methylation at CpG islands, shelves and shores we clustered the 11,323 variable regions from 3,825 CpG Islands, 4,821 CpG shelves and 2,677 CpG shores (Fig 4b). The resulting horizontal ordering of the samples shows a gradual change in DNA methylation from normal mucosa to adenocarcinoma. DMR clusters 1-4, containing >75% CpG shores and shelves, exhibit a general decrease in DNA methylation from normal mucosa to adenocarcinoma (Hypo-clusters), while clusters 5-8, consisting of >75% CpG islands, show an increase in DNA methylation from normal mucosa to adenocarcinoma (Hyper-clusters) (Fig 4c). Focusing on the chromatin state in Hyperclusters using public ChIP-seq data of four histone marks (H3K4me1, H3K4me3, H3K27ac and H3K27me3) in normal colon^34^ we found that >85% of the regions in the Hyper-clusters carry both an activating mark (H3K4me1) and a repressive mark (H3K27me3; Fig 4d) which is a hallmark of poised/bivalent genes^19^. Histone marks at the Hypo-clusters show weak ChIP-seq levels and thus their chromatin state could not be defined (Supplementary Fig 4b). Gene enrichment analysis of the genes associated with Hyper-clusters (Fig4 e; closest gene (<1000bp)) revealed a strong enrichment for general developmental programs such as regionalization, cell fate commitment and more. TCGA expression of the genes associated with the CpG islands/shelves and shores in each cluster (Supplementary Fig4c) shows a global decrease in expression from normal to tumor, with the decrease being most significant in clusters 6-7 that gain methylation and are enriched with CpG islands.

One example for a bivalent gene promoter can be seen at the IRF4 gene that has no observed accessibility at the gene promoter region while having very strong H3K4me1 and H3K27me3 signals (Fig 4f). The IRF4 promoter exhibits gradual increase in methylation level and its gene body region has a gradual decrease in methylation levels. These epigenetic modifications all occur with small and nonsignificant expression changes observable in TCGA data (Supplementary Fig 4d).

### Enhancer methylation show non-linear dynamics during CRC progression

The availability of ATAC-seq data for similar FAP samples enables us to follow the dynamics of DNA methylation and chromatin accessibility in detail. We examined the methylation changes at distal regulatory elements by integrating accessibility data defined using scATAC from a parallel dataset of benign-dysplasia-adenocarcinoma tumor progression in FAP patients^36^. We first filtered the accessible peaks by distance from CpG islands and transcription start site (>1 kb from both) and by the number of CpGs in each peak (>3 CpGs per peak; Supplementary Fig 4e). We then sorted the resulting 165,297 peaks based on the observed variability in the methylation levels across the samples and selected the top 42,593 that had >15% change in DNA methylation between groups for further evaluation. K-means (K=6) clustering of the methylation levels at these enhancer regions reveals a strong global change in methylation pattern between the groups (Fig 5a). Most of the variable enhancer regions present a non-monotonic methylation dynamic consisting of decreasing methylation with the progression from normal mucosa to benign and dysplastic states but elevated methylation in the adenocarcinoma state. Chromatin accessibility changes between the different disease states mirror the methylation changes in the same regions (Fig 5b). For example clusters 2 and 5 which gain methylation in tumor samples exhibit decreased DNA accessibility, while clusters 1,3,4 and 6 that lose DNA methylation gain DNA accessibility. Testing for enrichment of transcription factor motifs within each of the different clusters (Fig 5c) revealed similar groupings to the ones reported by analysis of single cell ATAC seq data^36^ with a very strong JUN:FOS (AP-1) enrichment at clusters that lose methylation (gain accessibility) and CDX, OLIG, GATA and NERUOD family transcription factors at clusters that gain methylation (lose accessibility). Specifically, cluster 5 shows a very strong enrichment for the HOX family transcription factors and also shows the highest level of hyper-methylation in adenocarcinoma samples.

**Figure 5:**
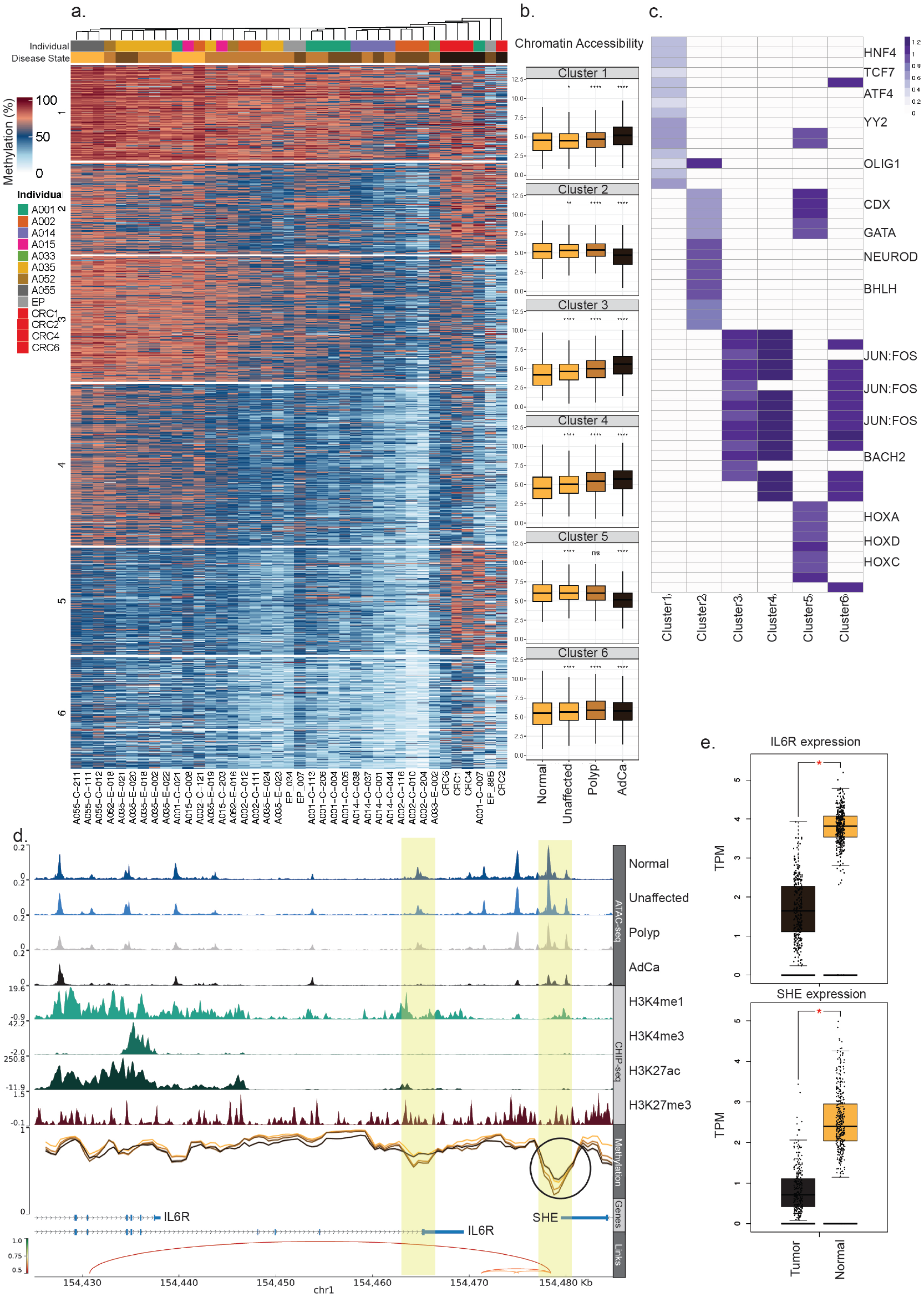
Strong changes in DNA methylation across thousands of colon regulatory elements. (a) K-means (K=6) clustering of 42,593 variable regulatory regions (defined by 500 bp-wide scATAC-seq peaks^35^) of all FAP samples colored as in Fig 4b. (b) scATAC-seq accessibility levels inpeaks found in clusters from panel a grouped and colored by pathological classification (x-axis; color code as in Fig 4b). (c) Transcription factor position weight matrix (PWM) enrichment in the 15 peaks with most significant p-values found in each cluster from panel a (TF with statistical significance of log10(p-value) >10 are shown). Purple color intensity codes for log2-enrichment of PWM over background (scATAC-seq peakset). (d) Genomic tracks of the *IL6R* gene loci as in Fig 4f where the lowest panel shows connection of enhancer regions to promoter regions. (e) TCGA RNA expression (y-axis) of *IL6R* and *SHE* at tumor vs normal COAD samples.

A typical example for an enhancer with nonmonotonic methylation dynamics is the *IL6R/SHE* enhancer (Fig 5d) which loses methylation in the progression from normal mucosa to the dysplastic state, but regains methylation in adenocarcinoma. Chromatin accessibility in these enhancer regions decreases with disease progression and the expression of associated genes (*IL6R* and *SHE)* has been shown to decrease in (CRC) tumor (TCGA; Fig 5e). These results demonstrate the complex choreography of regulatory events that occur during early CRC formation and progression.

## Discussion

DNA methylation has long been considered a marker for cancer progression, aging and disease state^37^. To date, many large scale landmark methylation studies balanced the scope and depth of their assays using arrays, RRBS or target capture methods^21,34,38^. Despite the ability of these assays to capture many biologically relevant changes along with DNA methylation levels, they capture only a fraction of genomic CpG sites that exhibit functionally meaningful changes in disease and aging. More recent studies have demonstrated that whole genome methylation sequencing (WGMS) indeed reveals many informative regions whose DNA methylation changes during aging, cancer progression and cell differentiation and provides a deeper understanding of biological mechanisms^39–44^.

Here, we present the utility of a novel sequencing technology (UG) that enables cost-efficient generation of WGMS data at scale. We validated the equivalence of methylation data generated on this platform using standard samples and criteria defined by the EpiQC consortium in terms of general sequencing criteria, conversion controls and genomic coverage. Furthermore, we demonstrate that detection of regions of differential methylation between two genome-in-a-bottle cell lines (HG001 and HG002) by UG are highly similar to existing reference data.

Comprehensive WGMS can unlock information on the function of genomic regions that have not been explored so far. For example, focusing on distal regulatory elements we measured changes in methylation state of 1,350,249 CpGs in over 165,000 distal peaks defined by scATAC-seq. Notably, only 63,363 CpGs (<5%) of these are represented at the Infinium MethylationEPIC array (850k array). Furthermore, we found that of the >15% of these CpGs (224,765 out of 1,350,249) that exhibit significant variability explored in our study, only <6% (12,320 out of 224,765) are covered by the array.

Familial adenomatous polyposis (FAP) patient samples offer a unique and valuable view of cancer progression. Although samples are collected at the same time point they cover multiple stages of tumor progression representing the malignancy trajectory and as such provide an exceptional resource for understanding early events that precede adenocarcinoma formation. Using UG sequencer we generated a unique dataset of WGMS at high coverage (>50X) on 44 samples taken from 15 patients. In this work we were able to detect millions of CpGs that change in the trajectory from normal mucosa through benign and dysplastic polyps to adenocarcinoma. Remarkably, hundreds of thousands of these CpGs become differentially methylated already in the benign and dysplastic states suggesting that many of the cancerous cellular transformations occur at this early stage. Although most of the differential changes that we detect reflect hypomethylation from the normal mucosa (144,284 hypo DMRs vs 6224 hyper DMR) only 10% of the changes in normal to tumor are observed in the early stages while a larger fraction (25%) of the normal mucosa to adenocarcinoma hyper-DMRs are already observed in the benign stage (Fig 3a,b). Comparison with TCGA data revealed that a larger fraction of the hypermethylated CpGs (70% as compared to 40% overlap in hypo-DMRs) are specific to colorectal tumors, suggesting a more tumor-specific behavior in hyper-methylation which occurs at tightly-regulated CpG islands. The gradual nature of methylation changes during CRC progression is observed in the increase in correlation with methylation changes in colorectal and rectal adenocarcinoma (Figs 3f,g), suggesting again that early occurring methylation changes are indeed part of the malignant transformation.

By integrating ATAC-seq and WGMS datasets we demonstrated the resemblance in these signals both in terms of correlated changes in specific regulatory elements as well as similar gene sets emerging from the cluster analysis (Fig 5c; Becker et al 2022^36^). These similarities highlight the fact that these two assays mirror similar underlying biological changes and to suggest that WGMS results can serve as proxy to the underlying chromatin state. Furthermore, by providing the methylation haplotypes of single molecules, WGMS provides further information on subpopulation of cells and how they can be distinguished from one another.

Individual sample clustering (Fig4b, Fig5a) based on WGMS levels revealed a gradual methylation change in pathologist-defined malignancy states, but also revealed clustering of individuals. Specifically, patients A014 and A035 form clusters which include multiple malignancy states. This observation may relate to the genotype of patient A014, which lacks the germline APC mutation^45^, and the polyps of patient A035 which present a sessile phenotype. These results mark the complexity of the DNA methylation phenotype and its specificity to individual patients and their health records. This emphasizes the need to combine multiple datasets of each individual patient.

In summary, this study demonstrates the utility of a novel sequencing platform in uncovering genome-scale information on the molecular mechanisms underlying the trajectory from normal mucosa to adenocarcinoma. Although analysis of additional samples is required to further characterize the physiological impact of these changes, we believe that the comprehensive dataset produced in this study will serve as a valuable resource for future studies and as a demonstration of the potential impact of this approach.

## Methods

### Patient Selection and Sample collection

Patient selection and sample collection were done as described in Horning et al^45^.

### NGS Library preparation

To generate whole genome DNA methylation libraries, approximately 200 ng of genomic DNA from each Genome in a Bottle (GIAB) reference sample (HG001, HG002, and HG005) (Coriell) and FAP samples was mechanically sheared to a fragment size of 200-300 bp (short insert) or 300-400 bp (long insert, EM-seq only) using Covaris. For EM-seq libraries, NEBNext® Ultra II DNA Library Prep reagents (NEB #E7645) were used to ligate 5mC-protected adapters. The NEBNext® Enzymatic Methyl-seq Conversion Module (NEB #E7125) was used to perform a two-step enzymatic conversion of non-methylated cytosines to uracils. For bisulfite-converted (Bisulfite-seq) libraries, fragmented gDNA was denatured and nonmethylated cytosines were converted to uracils using the EZ DNA Methylation-Gold Kit (Zymo D5005). After conversion, single stranded library prep was carried out using the xGen™ Methyl-Seq DNA Library Prep Kit (IDT #10009860). In the final PCR step for EM-seq and Bisulfite-seq libraries, Ultima Genomics (UG)-specific indexing primers were used.

For all EM-seq libraries we followed New England Biolabs (NEB) recommendations of ~200 bp insert size resulting in an average insert size of ~180 bp (Supplementary table 1).

### Ultima Genomics sequencing

Ultima Genomics sequencing was done as previously described in Almogy et al^24^

### Data processing for GIAB and FAP samples

Raw reads in FASTQ format were first trimmed for Illumina P5 adaptors and base quality using Cutadapt^46^ (- q 20,20 -e 0.2 -m 50 -g ACACGACGCTCTTCCGATCT), reads were then mapped to HG38 reference (containing pUC19 and Lambda genome) using BWA-Meth^47^ with default parameters. Output BAM file was then deduplicated and sorted using MarkDuplicatesSpark from GATK^48^. To call methylation levels we used both MethylDackel (https://github.com/dpryan79/MethylDackel) and WGBS-Tools (https://github.com/nloyfer/wgbs_tools). Mbias plots were produced using MethylDackel with default parameters.

## Analytical methods

### Differentially Methylated Regions (DMRs)

For DMR analysis we used Metilene^31^. Input CpG data for metilene was first filtered by coverage (>=5 in GIAB samples, >=10 in FAP samples) on all CpGs in the pairwise test. We ran metilene with the following flags -d 0 -m 5 -v 0.2, we then filtered the results by >15% methylation diff and by either qvalue < 0.05 or KS-test < 0.05.

### Cell type deconvolution

Prediction of cell type composition in our FAP and CRC data was done using the meth_atlas package (https://github.com/nloyfer/meth_atlas)^27^. To this end, WGMS data was reduced to 450 k array format.

### TCGA methylation and expression data

Processed methylation data from TCGA was obtained from the DNA Methylation Interactive Visualization Database (DNMIVD)^33^. Differential methylation was determined per CpG site and cancer type, and filtered to include methylation difference (tumor vs. normal) > 15%, adjusted p-value < 0.05. As TCGA data is of 450 k array, matched CpGs were extracted from our WGMS. Sites common to all 21 cancer types and our data include 339,977 CpGs. Gene expression data of the COAD dataset was downloaded as “htseq-counts” from UCSC Xena browser.

### K-means clustering

K-Means clustering of average methylation across samples was done using Kmeans++ algorithm. Clustered data was then sorted using hclust both on the samples (horizontal) and clusters (vertical).

### Transcription factor enrichment

To generate motif match matrices, motifs from the curated, high-confidence JASPAR2020 vertebrate core database were obtained^49^. We called significant motif matches in peaks, or subsets of peaks, using JASPAR motif position weight matrices and the function “matchMotifs” from the “motifmatchr” and “chromVAR” packages, resulting in a binary peaks-by-matches matrix. To compute enrichments, we defined foreground (specific cluster) and background peak sets (all scATAC-seq peaks), then performed a two sided Fisher’s Exact test for the over representation of each motif in the foreground set. For K-means cluster motif enrichments, we aggregated all peaks from a given K-means cluster and tallied the total number of matches for each motif in the cluster, and we used all of the other clusters as the background

## Data Availability

The datasets used and/or analyzed during the current study are available from the corresponding author upon request.

## Ethics declarations

G.K, T.C., E.M., S.G., S.B., D.Lebanony., F.O, N.I., A.J., D.L. and Z.S. are employees and shareholders of Ultima Genomics.

M.P.S. is a cofounder and scientific advisor for Personalis, Qbio, January.ai, Filtricine, Mirvie, Protos and an advisor for Genapsys.

## Author contributions

### Conceptualization

H.L., J.F., M.P.S.

### Sample collection and processing

H.L., T.C., A.K., A.H., C.R.H., Y.Z., E.D.E., S.N., A.W., E.M., S.G., S.B., D.Lebanony.

### Ultima sequencing

F.O., N.I.

### Computational analysis

H.L., G.K., A.J., Z.S.

### Supervision

A.J., D.L., Z.S., M.P.S.

### Writing

H.L., G.K., A.J., D.L., Z.S., M.P.S.

