## Supplemental Figures 1-4 for "Ultra high-throughput whole-genome methylation sequencing reveals trajectories in precancerous polyps to early colorectal adenocarcinoma"

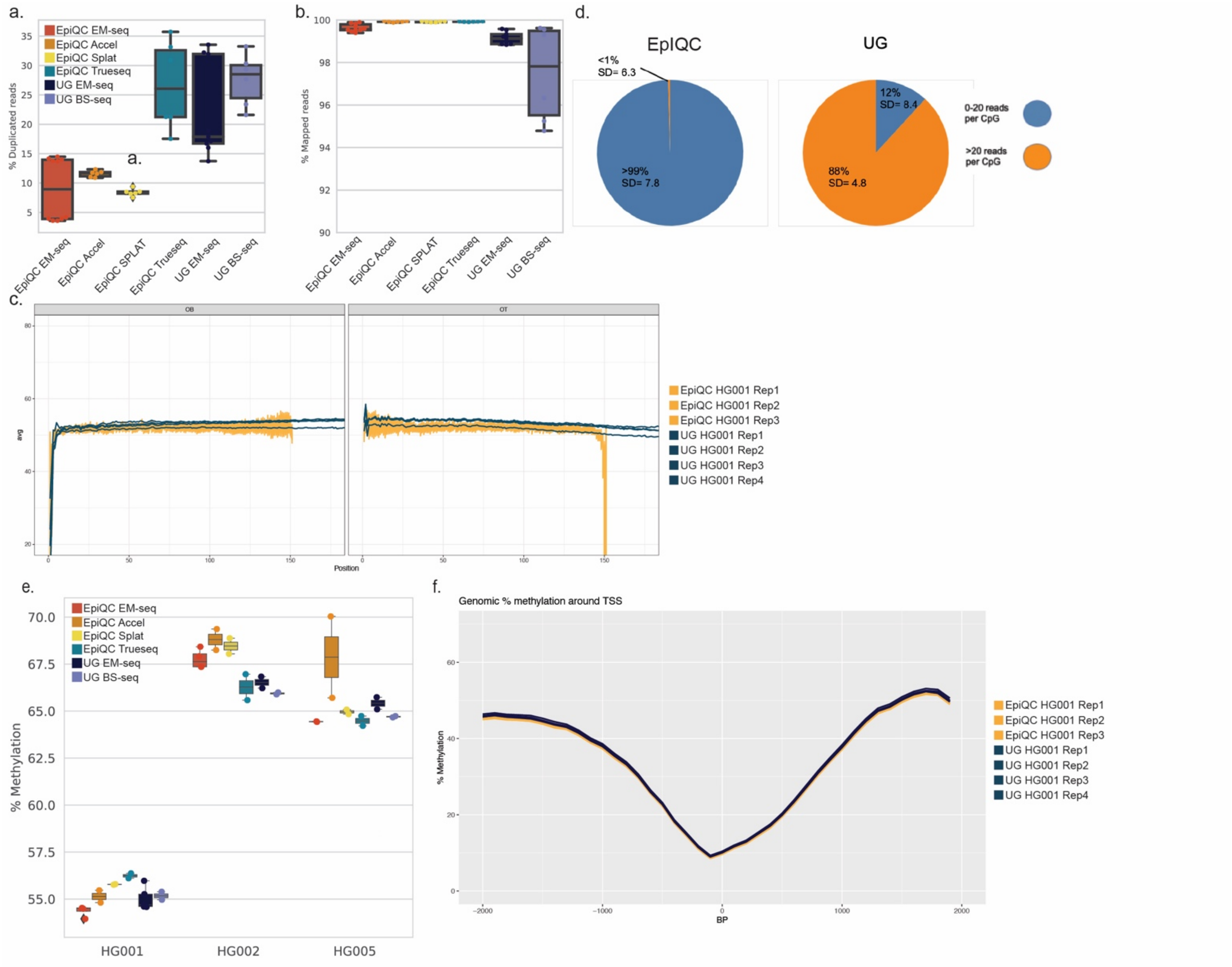


**Supplementary Figure 1**: Quality controls for WGMS of genome in a bottle cell lines. (a) Percent of duplicated reads. (b) Percent of mapped reads. (c) M-bias plots, showing percent methylated CpGs along the read. Shown are EM-seq samples of EpiQC and UG. (d) Pie charts showing the percent of highly covered CpGs (>20 reads) and lowly covered CpGs (<20 reads) in two EM-seq samples of EpiQC (left) and of UG (right). Standard deviation of methylation levels between replicates is shown per coverage section. (e) Average genomic methylation level of EpiQC and UG samples, per genome. Each point is a single sample. (f) Methylation level around transcription start sites (TSS). Shown are EM-seq samples of the two platforms.

a.

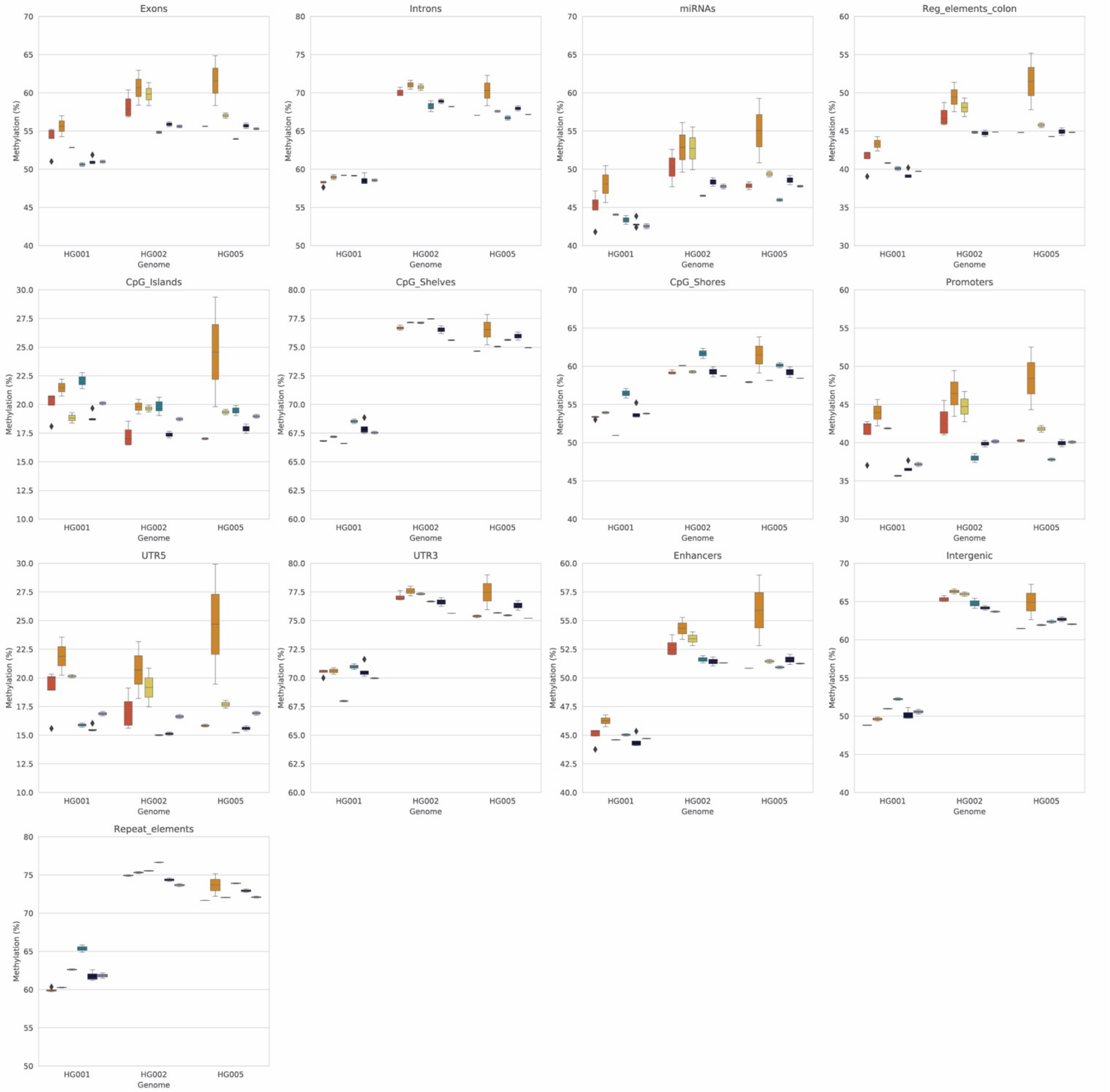


**Supplementary Figure 2:** (a) Methylation at different genomic features, presented for EpiQC and UG samples.

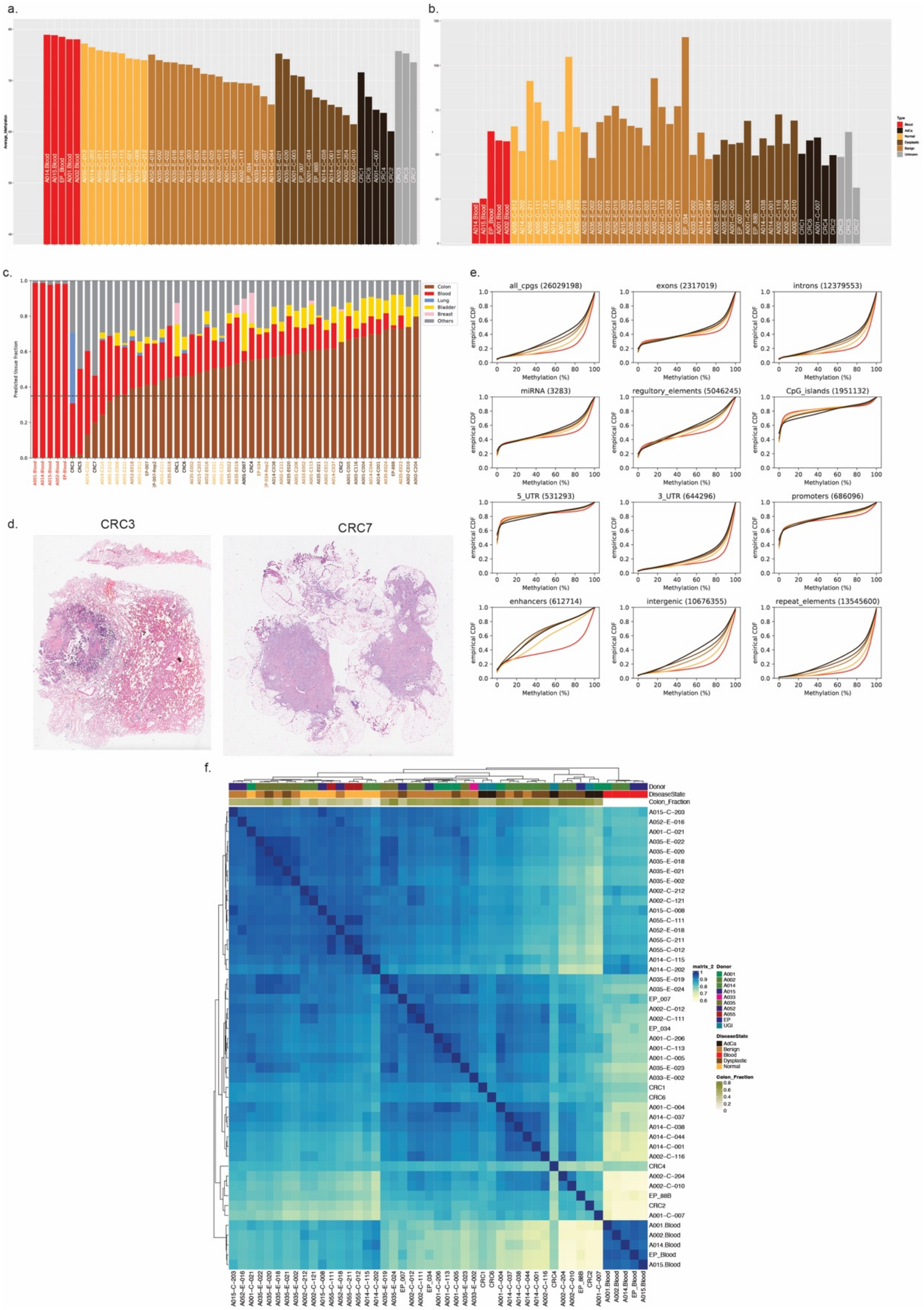


**Supplementary Figure** 3: Quality controls for FAP samples. (a) Average genomic methylation level of FAP samples. Samples in gray were filtered out of downstream analysis. (b) Average CpG coverage per sample. (c) Cell type composition. Shown is the predicted fraction of each cell type per sample, using methylation based-cell type deconvolution analysis. (d) H&E slides of two CRC samples that were filtered out from the analysis. (e) Cumulative distribution function graphs of average methylation levels per genomic features. Shown is the mean of each pathologist-defined group. (f) Correlation between samples. Shown is the Pearson correlation coefficient between samples based on all CpGs with minimal coverage of 10 reads.

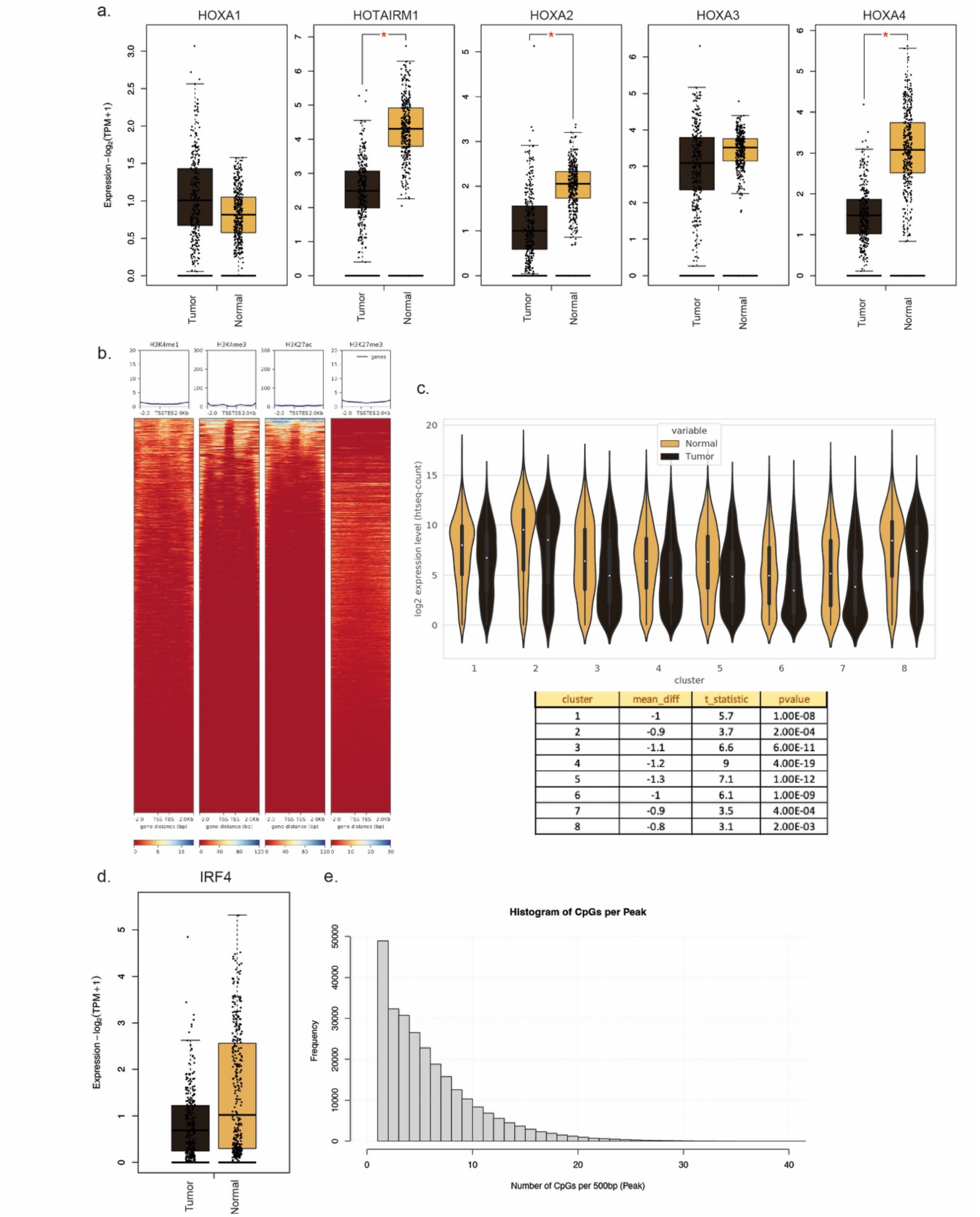


| cluster | mean_diff | t_statistic | pvalue |
|---|---|---|---|
| 1 | -1 | 5.7 | 1.00E-08 |
| 2 | -0.9 | 3.7 | 2.00E-04 |
| 3 | -1.1 | 6.6 | 6.00E-11 |
| 4 | -1.2 | 9 | 4.00E-19 |
| 5 | -1.3 | 7.1 | 1.00E-12 |
| 6 | -1 | 6.1 | 1.00E-09 |
| 7 | -0.9 | 3.5 | 4.00E-04 |
| 8 | -0.8 | 3.1 | 2.00E-03 |

**Supplementary Figure 4:** CpG Islands and Distal regulatory regions. (a) TCGA average gene expression of genes found in the *HOX* locus (b) Histone marks around CpG Islands/Shelves and Shores found at clusters 1-4 from Figure 4b. (c) TCGA gene expression of genes associated with CpG Islands/Shelves and Shores (nearest gene, up to 1000bp from TSS) found at clusters in Figure 4b. (d) *IRF4* TCGA gene expression. (e) Number of CpGs found within all scATAC-seq peaks.
